# Rhizosphere bacterial community composition depends on plant diversity legacy in soil and plant species identity

**DOI:** 10.1101/287235

**Authors:** Marc W. Schmid, Terhi Hahl, Sofia J. van Moorsel, Cameron Wagg, Gerlinde B. De Deyn, Bernhard Schmid

## Abstract

Soil microbes are known to be involved in a number of essential ecosystem processes such as nutrient cycling, plant productivity and the maintenance of plant species diversity. However, how plant species diversity and identity affect soil microbial diversity and community composition is largely unknown. We tested whether, over the course of 11 years, distinct soil bacterial communities developed under plant monocultures and mixtures, and if over this timeframe plants with a monoculture or mixture history changed in the microbial communities they associated with. For eight species, we grew offspring of plants that had been grown for 11 years in the same monocultures or mixtures (monoculture- or mixture-type plants) in pots inoculated with microbes extracted from the monoculture and mixture soils. After five months of growth in the glasshouse, we collected rhizosphere soil from each plant and used 16S-rRNA gene sequencing to determine the community composition and diversity of the bacterial communities. Microbial community structure in the plant rhizosphere was primarily determined by soil legacy (monoculture vs. mixture soil) and by plant species identity, but not by plant legacy (monoculture- vs. mixture-type plants). In seven out of the eight plant species bacterial abundance was larger when inoculated with microbes from mixture soil. We conclude that plant diversity can strongly affect belowground community composition and diversity, feeding back to the assemblage of rhizosphere microbial communities in newly establishing plants. Thereby our work demonstrates that concerns for plant biodiversity loss are also concerns for soil biodiversity loss.

## 1 INTRODUCTION

Soil microbes play an essential role in a number of ecosystem processes including decomposition, nutrient cycling, plant productivity and the maintenance of plant species diversity (Ahemad & Kibret, 2014; Bever, Mangan, & Alexander, 2015; Wagg, Bender, Widmer, & van der Heijden, 2014). Loss of soil biodiversity reduces ecosystem functioning as trophic networks collapse (Gosling, Hodge, Goodlass, & Bending, 2006; Wagg et al., 2014) and plants are increasingly exposed to specialized soil-borne pathogens from which they are normally protected by the large biodiversity of other soil organisms (Ahemad & Kibret, 2014; Eisenhauer, Reich, & Scheu, 2012; van der Putten et al., 2013). The role of potential feedbacks of plant diversity on soil diversity and rhizosphere community assemblages, however, is largely unexplored (Dassen et al., 2017).

Soil microbes form tight associations with plants (Bulgarelli, Schlaeppi, Spaepen, van Themaat, & Schulze-Lefert, 2013) and key mutualistic to antagonistic interactions between plants and soil biota take place in the rhizosphere, defined by a narrow zone of soil surrounding plant roots (Whipps, 2001). The composition of microbial communities in the rhizosphere is mostly determined by local biotic and abiotic conditions (van der Putten et al., 2013), of which the composition of the local plant community may contribute to shaping these conditions (Bardgett & Wardle, 2003). Plants can initiate large compositional changes in rhizosphere microbiomes (Dakora & Phillips, 2002; Latz, Eisenhauer, Rall, Scheu, & Jousset, 2016) and the conditions in the rhizosphere can vary strongly between plant species (Berg & Smalla, 2009; Eisenhauer et al., 2017; Latz et al., 2016). Considering the unique influence of different plant species on shaping the soil microbiome the loss of plant species likely results in a loss soil microbial biodiversity (Broughton & Gross, 2000; Garbeva, Postma, van Veen, & van Elsas, 2006; Hooper et al., 2000; Schlatter, Bakker, Bradeen, & Kinkel, 2015). However, it is unclear to which extent plant species richness, or plant species identity, affect soil microbial diversity and composition. A recent study suggests that the role of differences between plant functional types may be larger than that of plant species richness *per se* on the composition of the soil microbiome (Dassen et al., 2017). In addition, higher plant diversity often increases plant aboveground biomass (Balvanera et al., 2006; Reich et al., 2012) that can sustain a larger amount of soil bacteria and fungi (De Deyn, Quirk, & Bardgett, 2011).

Consequences of reduced plant biodiversity for soil biodiversity can be studied in long-term biodiversity experiments (Eisenhauer et al., 2011; Roscher et al., 2013; Tilman, Reich, & Knops, 2006; Zuppinger-Dingley, Flynn, De Deyn, Petermann, & Schmid, 2016). These experiments can also be used as selection experiments (Cardinale et al., 2012; Eisenhauer et al., 2016). Over time, different selection pressures in experimental monocultures vs. mixtures can result in pools of plant genotypes adapted to monocultures vs. mixtures (van Moorsel et al., 2018; Zuppinger-Dingley et al., 2014; Zuppinger-Dingley, Flynn, Brandl, & Schmid, 2015; Zuppinger-Dingley et al., 2016). Here we refer to the monoculture- and mixture-selected plant genotypes as monoculture- and mixture-type plants, respectively. The formation of such plant sub-types may occur via a sorting-out from standing genetic variation (Fakheran et al., 2010). Monoculture- and mixture-type plants have been shown to be distinguishable from each other after 8–11 years of selection based on plant performance and functional trait variation (van Moorsel, Schmid, Hahl, Zuppinger-Dingley, & Schmid, 2018; Zuppinger-Dingley et al., 2014). In addition, Zuppinger-Dingley et al. (2016) found that after eight years of co-development of plant and soil communities, feedbacks of microbes from monoculture soil were positive for monoculture-type plants, but negative for mixture-type plants of the same species. The authors suggested that an accumulation of specialized pathogens in monocultures, and their dilution in mixtures, could create differential selection pressures on plants (Zuppinger-Dingley et al., 2016). These studies, however, were unable to investigate the community composition of soil microbes and how they may respond to plant diversity.

Here we ask whether differences in plant species diversity and differences in plant history can alter soil microbial communities colonizing the rhizosphere of different plant species, resulting in a legacy effect on the plant species rhizosphere microbiome associations. We tested whether, over the course of 11 years, i) distinct soil microbial communities developed under plant monocultures (monoculture soil) and mixtures (mixture soil) and ii) plant history (monoculture- vs. mixture-type plants) alters the assembly of rhizosphere communities in a large field biodiversity experiment in Jena, Germany (the Jena Experiment, Roscher et al., 2004). We grew monoculture- or mixture-type plants of eight species belonging to four different functional groups as single individuals in pots inoculated with microbes extracted from the monoculture and mixture soils. We assessed the influence of soil legacy (monoculture vs. mixture soil), plant species identity and plant legacy (monoculture- vs. mixture-type plants) on the composition of the soil microbiome in the rhizosphere of the potted test plants. We hypothesize (*i*) that microbiomes obtained from mixture soil are more diverse and differ in composition from microbiomes obtained from monoculture soil. Furthermore, we expect (*ii*) that plant species differ in their microbiomes. Finally, we hypothesize (*iii*) that monoculture- and mixture-type plants obtain different microbiomes when inoculated with microbes from the same soil if plant-soil microbiome associations have been co-selected through their shared history under field conditions. We used 16S-rRNA gene sequencing to determine the community structure and diversity of the bacterial communities. Because 16S amplicon read frequencies cannot be used to compare abundances between species (Edgar 2017), we did not analyze variation in abundance-weighted diversity metrics such as evenness, dominance or Shannon diversity (Magurran 2004). Instead, to characterize the overall impact of soil legacy, plant species identity and plant legacy on the community structure of rhizosphere bacteria, we analyzed the variation in the total number of detected operational taxonomic units (OTUs) and the variation in abundances within bacterial OTUs across treatments.

## 2 METHODS

### 2.1 Plant species

We used eight common European grassland plant species previously classified into different functional groups (Roscher et al., 2004): one grass (*Festuca rubra* L.), three small herbs (*Plantago lanceolata* L., *Prunella vulgaris* L., and *Veronica chamaedrys* L.), two tall herbs (*Galium mollugo* L. and *Geranium pratense* L.) and two legumes (*Lathyrus pratensis* L. and *Onobrychis viciifolia* Skop.). The studied species had undergone 11 years of selection in either plant monocultures (monoculture-type plants) or species mixtures (mixture-type plants) from 2002–2014 (see Fig. 1).

**Figure 1.**
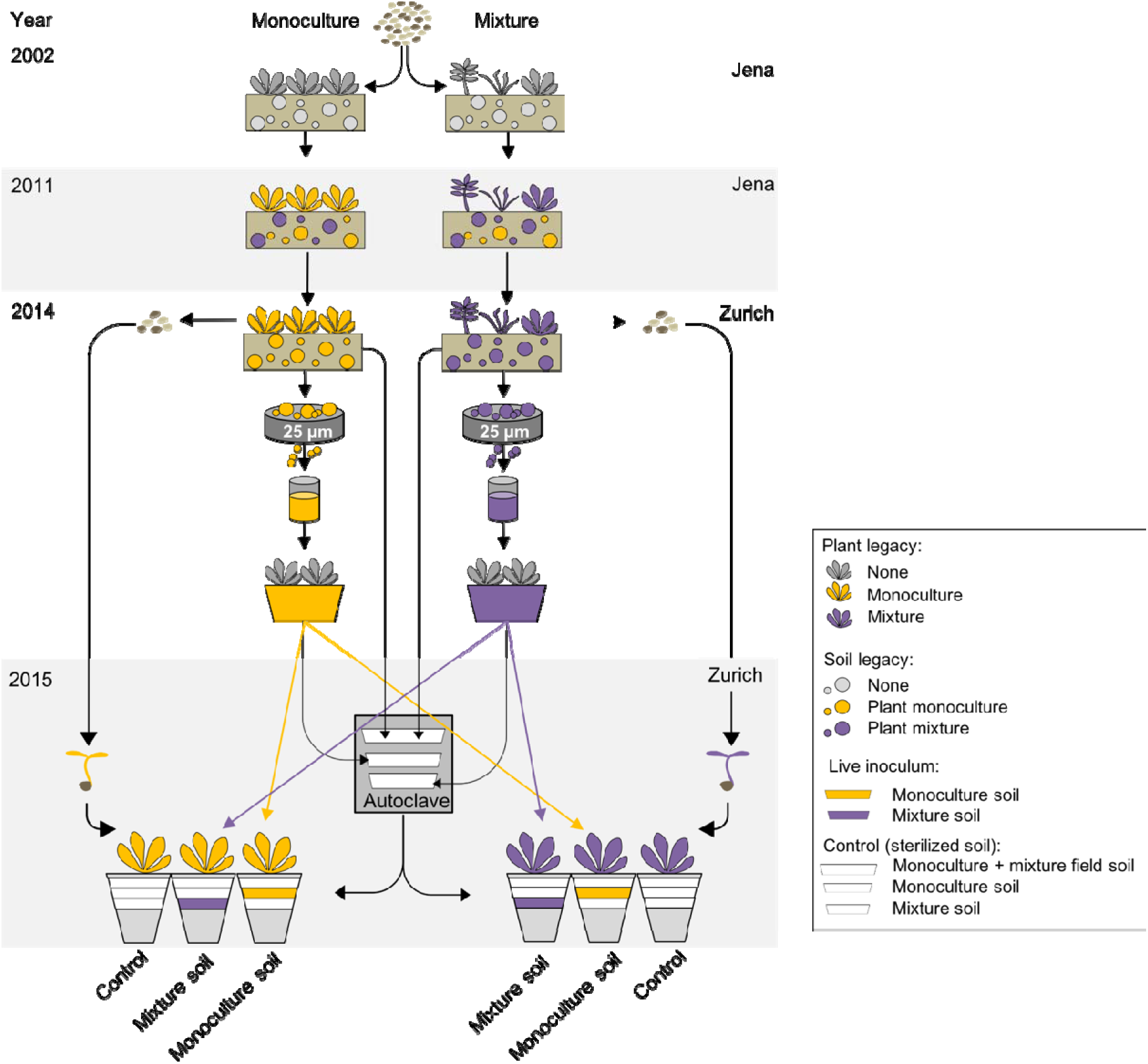
Experimental design. Plant monocultures and mixtures were sown in the Jena Experiment in 2002 and maintained until 2010. In 2010 the soil of the plots was pooled and placed back to the same locations. In spring 2011, plant seedlings were planted in the mixed soil in identical species composition as their parents. The soil communities were allowed to re-assemble with their original plant communities for three more years from 2011–2014. In spring 2014, rhizosphere soils from eight plant species were collected and the plants were used for a second controlled seed production. Soil microorganisms smaller than 25 µm in diameter (i.e. excluding mycorrhizal spores) were then isolated. The isolated micro-organisms were allowed to accumulate in trap-cultures for five months with neutral trap plants for each of the eight plant species. To create soil treatments for the subsequent pot experiment in the glasshouse, we filled pots with sterile soil (in grey) and added live inoculum of either microbes isolated from plants grown in monoculture (monoculture soil, indicated in yellow) or microbes isolated from plants grown in mixture (mixture soil, indicated in purple). To standardize the nutrient composition between pots, we added an 0.8 dl autoclaved counterpart of the remaining inocula to each pot (indicated in white, for details see Methods and Hahl, 2017). The control soil treatment received the same amount of each inoculum, but all inocula were autoclaved. Finally, we added 1 dl of the gamma-radiated sand-soil mixture to avoid cross-contamination of the live soil inocula between pots (indicated in grey). Then a single monoculture- or mixture-type plant (drawn in yellow or purple, respectively), germinated from the seeds of second controlled seed production, was planted to each pot.

### 2.2 Producing plants and soils with monoculture vs. mixture legacy

Plant communities of 48 plots (12 monocultures, 12 two-species mixtures, 12 four-species mixtures and 12 eight-species mixtures) of a field biodiversity experiment in Jena, Germany (the Jena Experiment, see Roscher et al., 2004), were collected as cuttings in spring 2010, after eight years of growth in their respective plant communities. These cuttings were transplanted in identical plant composition to an experimental garden in Zurich, Switzerland, for the first controlled sexual reproduction among “co-selected” plants (for details see Zuppinger-Dingley et al., 2014). In addition, the top 30 cm of soil of the 48 plots was pooled, mixed and returned to the excavated locations in the Jena Experiment. In spring 2011, the seedlings produced from the seeds of the first controlled sexual reproduction were transplanted back to this mixed soil in the same plots of the Jena Experiment from where the parents had originally been collected and in the same community composition as the parents had been established. These plant communities were maintained for three years until 2014 to allow them to become associated again with their own microbial communities and continue the selection treatments in their respective communities (Fig. 1, upper part).

In March 2014, plant communities including rhizosphere soil of the re-established plots in the Jena Experiment were transplanted to plots in the experimental garden in Zurich for the second controlled sexual reproduction. The plots had been filled with 30 cm of soil (1:1 mixture of garden compost and agricultural soil, pH 7.4, Gartenhumus, RICOTER Erdaufbereitung AG, Aarberg, Switzerland) and fenced with netting to minimize cross-pollination with plants outside the plots. Seeds of monoculture-type plants were collected from monoculture plots and seeds of mixture-type plants form four- or eight-species mixture plots of 1 x 1 m in the experimental garden. After collection, the seeds of the eight plant species were stored at +4°C for at least two months. This plant material was then used in the pot experiment in the glasshouse described below.

### 2.3 Soil inoculum preparation

In March 2014, rhizosphere soil samples attached to the roots of the plants that we transported to Zurich for the second sexual reproduction were collected and stored at 4° C. The monoculture soils came from the eight plant monoculture plots and the mixture soils came from seven different eight-species plant mixture plots (one for each species except for *G. mollugo* and *O. viciifolia* whose rhizosphere soil samples came from the same mixture plot) in the Jena Experiment. Microbial communities of the sampled rhizosphere soil were isolated and propagated and subsequently used in the pot experiment in the glasshouse. To isolate the microbial communities, we produced a microbial wash by passing 500 ml of deionized water and 25 g of rhizosphere soil through a series of sieves with the smallest mesh size of 25 µm (Koide & Li, 1989, Wagg et al., 2014). To propagate the isolated microbes, we established trap cultures in two replicates for the eight plant species but using seeds without the above-mentioned legacies (seeds from Rieger-Hofmann GmbH, Blaufelden-Raboldshausen, Germany). The trap cultures consisted of 2 L pots filled with an autoclaved (120° C for 99 min) sand-soil mixture (4:1) and planted with several trap plant individuals (surface-sterilized seeds pre-germinated on 1 % water-agar) per species and pot (Fig. 1). At the same time as the planting each trap culture received 250 ml of microbial wash. After five months of growth in the glasshouse we pooled the soils of the replicated trap cultures per plant species and soil legacy (monoculture or mixture soil). Trap plant roots were cut into 3–5 cm fragments and used together with the soil as inoculum in the pot experiment as described below.

### 2.4 Setup of the pot experiment in the glasshouse

We filled 1-L pots with 5.6 dl of gamma-radiated (27–53 kGy) 1:1 (w/w) sand/soil mixture (RICOTER Erdaufbereitung AG) and added 0.8 dl inoculum from three sources (see Fig. 1). The soil-legacy treatments were thus a) control (no live inoculum), b) microbes isolated from soil under plant monocultures (monoculture soil) and c) microbes isolated from soil under plant mixtures (mixture soil). Inoculum was prepared in the following way: first, monoculture-soil treatments (b) received 0.8 dl of live inoculum from trap cultures of monoculture soil and mixture-soil treatments (c) received 0.8 dl of live inoculum from trap cultures of mixture soil. Second, monoculture-soil treatments (b) received 0.8 dl of autoclaved (99 min at 120° C) inoculum from trap cultures of mixture soil, mixture-soil treatments received 0.8 dl of autoclaved (99 min at 120° C) inoculum from cultures of monoculture soil and control soil treatments (a) received both of these autoclaved inocula. Third, we added 0.8 dl of autoclaved field soil (99 min at 120° C; this was for comparison with a forth treatment, which differed from the control by receiving live field soil but which is not included in the analyses presented here, see Hahl, (2017) and 1 dl of the gamma-radiated sand-soil mixture to avoid cross-contamination of the live soil inocula between pots.

Seeds were surface-sterilized and germinated for two to four weeks (depending on pre-tested germination times of each species) prior to the pot experiment on 1 % water-agar. One pre-germinated monoculture- or mixture-type plant seedling of one of the eight test species was planted in each pot. The experiment included in total three soil-legacy treatments (monoculture and mixture soil and control), eight plant species and two plant-legacy treatments (monoculture- and mixture-type plants). The design of the experiment was a full 3 x 8 x 2 factorial. Each treatment combination was replicated seven times resulting in 336 pots which we randomly arranged within seven experimental bocks in the glasshouse. After 19–23 weeks of plant growth, we collected rhizosphere soil samples from each pot containing a live plant and stored the samples at –80° C.

### 2.5 Library preparation and sequencing

DNA was isolated from 500 mg of rhizosphere soil using the FastDNA SPIN Kit for Soil (MP Biomedicals, Illkirch-Graffenstaden, France) following the manufacturer’s instructions. Samples from a subset of 150 plants were chosen for the molecular analyses (Table S1). We carried out targeted PCR in duplicates to amplify the variable region V4 of the prokaryotic ribosomal RNA gene using primers 515f (GTGCCAGCMGCCGCGGTAA) combined with 5’ Illumina adapter, forward primer pad, and forward primer linker and barcoded 806r (GGACTACHVGGGTWTCTAAT) combined with Illumina 3’ adapter, Golay barcode, reverse primer pad, and reverse primer linker (Table S2, Bates et al., 2011). The PCR conditions for the amplification of the V4 region consisted of an initial denaturation at 94° C for 3 min, 30 cycles of denaturation at 94° C for 30 s, an annealing at 50° C for 30 s, and an elongation at 72° C for 1 min followed by a final elongation at 72° C for 10 min. The PCR products were purified using Agencourt AMPure XP magnetic beads (Beckman Coulter, Brea, USA). The amplicon concentrations were measured with the Fragment Analyzer and the Standard Sensitivity NGS Fragment Analysis kit (Advanced Analytical Technologies, Inc., Heidelberg, Germany). 60 ng of each sample were pooled and paired-end sequenced (2 x 300 bp) on the Illumina MiSeq 300 system (Beijing Genomics Institute, Bejing, China). Short-reads were deposited at SRA (accession number SRP105254).

**Table 1.** Analysis of variance of microbial richness (number of OTUs). Contrasts among soil-legacy treatments and their interactions are indented and printed in italics. Significant *P*-values are highlighted in bold.

| Source of variation | Df | Species richness |  |
| --- | --- | --- | --- |
|  |  | %-SS | *P* |
| Soil-legacy treatment (St) | 2 | 30.9 | **<0.001** |
| *Control vs. microbial soil (C)* | 1 | 30.6 | **<0.001** |
| *Monoculture vs. Mixture soil (M)* | 1 | 0.3 | 0.161 |
| Species (Sp) | 7 | 5.6 | **<0.001** |
| Plant-legacy treatment (Ph) | 1 | 0.0 | 0.594 |
| St × Sp | 14 | 5.6 | **0.003** |
| *C × Sp* | 7 | 3.2 | **0.007** |
| *M × Sp* | 7 | 2.4 | **0.039** |
| Sp × Ph | 7 | 1.1 | 0.454 |
| St × Ph | 2 | 0.2 | 0.499 |
| *C × Ph* | 1 | 0.1 | 0.367 |
| *M × Ph* | 1 | 0.1 | 0.448 |
| St × Sp × Ph | 14 | 2.4 | 0.381 |
| *C × Sp × Ph* | 7 | 1.2 | 0.370 |
| *M × Sp × Ph* | 7 | 1.2 | 0.388 |
| Residuals | 98 | 15.1 |  |

*Notes:* Df, degrees of freedom; %-SS, proportion of total sum of squares; *P*, error probability

**Table 2.** The number of OTUs exhibiting significant differential abundance in any of the contrasts tested in this study (FDR <= 0.01 and abs(logFC) >= 1; see Methods).

|  | All | *F. rubra* | *G. mollugo* | *G. pratense* | *L. pratense* | *O. viciifolia* | *P. lanceolata* | *P. vulgaris* | *V. chamaedrys* | Total (n) |
| --- | --- | --- | --- | --- | --- | --- | --- | --- | --- | --- |
| 1. PH: mixture- vs monoculture-type plants | 1 (0/1) | 5 (4/1) | 6 (5/1) | 0 | 3 (1/2) | 1 (1/0) | 6 (6/0) | 11 (4/7) | 1 (0/1) | 32 |
| a) Within control soil | 12 (6/6) | - | - | - | - | - | - | - | - | 120 |
| b) Within monoculture soil | 0 | 0 | 2 (2/0) | 0 | 2 (1/1) | 0 | 0 | 5 (0/5) | 0 | 9 |
| c) Within mixture soil | 0 | 0 | 0 | 0 | 0 | 0 | 1 (1/0) | 2 (1/1) | 0 | 3 |
| 2. SO: control vs. microbial soils | 972 (156/816) | 321 (76/245) | 267 (92/175) | 136 (29/107) | 171 (46/125) | 339 (37/302) | 295 (70/225) | 296 (77/219) | 366 (53/313) | 1026 |
| 3. SO: mixture vs. monoculture soil | 137 (105/32) | 136 (108/28) | 135 (85/50) | 212 (68/144) | 154 (112/42) | 103 (77/26) | 126 (69/57) | 241 (144/97) | 133 (68/65) | 844 |
| 4. Plant species: one vs. all others | - | 498 (192/306) | 129 (84/45) | 236 (167/69) | 221 (162/59) | 264 (148/116) | 290 (133/157) | 299 (118/181) | 500 (422/78) | 1287 |
| a) Within control soil | - | 96 (61/35) | 40 (35/5) | 54 (50/4) | 56 (54/2) | 35 (24/11) | 69 (42/27) | 63 (49/14) | 77 (69/8) | 388 |
| b) Within monoculture soil | - | 463 (182/281) | 218 (146/72) | 423 (290/133) | 157 (101/56) | 325 (194/131) | 302 (178/124) | 346 (173/173) | 593 (497/96) | 1465 |
| c) Within mixture soil | - | 346 (178/168) | 223 (130/93) | 286 (142/144) | 393 (285/108) | 281 (208/73) | 337 (163/174) | 402 (198/204) | 454 (393/61) | 1426 |

*Note:* Numbers in parentheses refer to the number of OTUs having higher abundance in the first and the second group of the contrast, respectively. Contrasts included either all species or each species individually (columns). The contrast comparing the different plant legacies (PH) was tested across the entire data set and within the specific soil-legacy treatments. Considering that plant legacy had a
weak effect on the composition of the microbiomes, the other contrasts comparing the different soil-legacy treatments (SO) were only tested across the entire data set and within the individual plant species but not within the specific plant-legacy treatments. Likewise, the contrast comparing one specific plant species with all others (SP) was only tested across the entire data set and within the specific soil-legacy treatments. Because there were only three samples from the control soil per species (i.e. one with monoculture- and two with mixture-type plants or *vice versa*), the contrast 1.a) was only tested across all species

### 2.6 Identification and annotation of OTUs

Operational taxonomic units (OTUs) were generated with UPARSE (version 8.1.1861, Edgar, 2013) following the example and the tutorial given for paired-end Illumina data (drive5.com/uparse/). Reads were first quality-checked with FastQC (bioinformatics.babraham.ac.uk/projects/fastqc). Following removal of primer sequences (Table S2) and low-quality bases with Trimmomatic (version 0.33 with the parameters ILLUMINACLIP:primerSeqs:2:30:10:8:1 SLIDINGWINDOW:5:15 MINLEN:100, (Bolger, Lohse, & Usadel, 2014)), paired-end reads were merged and filtered using usearch (with the parameters -fastq_maxdiffs 25 -fastq_maxdiffpct 10 for merging and - fastq_trunclen 250 -fastq_maxee 0.25 for filtering, Edgar, 2013). Duplicated sequences were then removed with fqtrim (version 0.9.4, Pertea, 2009). The remaining sequences were clustered with usearch (with the parameter -minsize 2, Edgar, 2013, 2017) to obtain 10’205 OTU sequences (Supplemental File S1). OTU sequences were annotated with the taxonomy data from SILVA (Quast et al., 2013) using SINA with a minimal similarity of 90 % and the 10 nearest neighbors (www.arb-silva.de/aligner, Yilmaz et al., 2014, Table S3). OTU abundances were finally obtained by counting the number of sequences (merged and filtered) matching to the OTU sequences (usearch with the parameters - usearch_global -strand plus -id 0.97, Edgar, 2013, Table S4). Three samples (Sample77, Sample265, and Sample364) were removed from all further analysis because they had very low counts (6, 12, and 1 counts in total). OTUs annotated as chloroplast were removed to avoid a potential bias caused by plant DNA. To avoid sequencing artifacts, OTU sequences with less than 50 counts in total or with counts in less than five samples were removed from all further analyses (4’339 OTUs remained after this filter).

### 2.7 Data normalization and identification of differentially abundant OTUs

Variation in OTU relative abundance was analyzed with a generalized linear model in R with the package DESeq2 (version 1.14.1, Love, Huber, & Anders, 2014) according to the factorial design with the three explanatory factors soil legacy (control, monoculture soil and mixture soil), plant species identity (*F. rubra*, *G. mollugo*, *G. pratense*, *L. pratensis*, *O. viciifolia*, *P. lanceolata*, *P. vulgaris*, and *V. chamaedrys*) and plant legacy (monoculture- and mixture-type plants). All individual factor combinations were coded as a unique level of a combined single factor (Table S1). Specific conditions were then compared with linear contrasts (Neter & Wassermann 1974). The four main contrasts compared (1) the two different plant legacies, (2) the control soil and the microbial soils, (3) the two different soil legacies of the microbial soils, and (4) each plant species to all other plant species. To test for interactions, each contrast was tested across the entire data set and within the individual soil-legacy treatments or different plant species. Contrasts (2), (3), and (4) were not tested separately within the two plant-legacy treatments because these only had weak effects on the composition of the microbiomes. Within each comparison, *P*-values were adjusted for multiple testing (Benjamini-Hochberg), and OTUs with an adjusted *P*-value (false discovery rate, FDR) below 0.01 and a minimal log2 fold-change (i.e., the difference between the log2 transformed, normalized OTU counts) of 1 were considered to be differentially abundant (Table S5). Normalized OTU counts were calculated accordingly with DESeq2 and log2(x+1)-transformed to obtain the normalized OTU abundances. Sequencing data were not rarefied (McMurdie & Holmes, 2014).

### 2.8 Enrichment and depletion of microbial phyla

To test for enrichment/depletion of microbial phyla occurrences in a given set of OTUs (e.g., OTUs with significant difference in abundance between monoculture and mixture soils), we constructed for each phylum a contingency table with the within/outside phyla counts for the given set of OTUs and all OTUs passing the filter. We then tested for significance with Fisher’s exact test. *P*-values were adjusted for multiple testing (Benjamini-Hochberg), and phyla with an adjusted *P*-value (false discovery rate, FDR) below 0.05 were considered to be significantly enriched/depleted (Table S7).

## 3 RESULTS

Amplification of 16S rRNA gene fragments yielded an initial set of 10’205 operational taxonomic units (OTUs, Supplemental File S1, Tables S3 and S4). After removing OTUs with similarity to chloroplast sequences (87 OTUs) and low-abundance OTUs (5’780 OTUs), 4’339 OTUs remained. Of these, 3’975 and 41 were classified as bacteria and archaea, respectively (194 remained unclassified). Within the bacterial domain, the ten most abundant phyla accounted for 86.7 % of all OTUs (Table S6); they were *Proteobacteria* (35.4 %), *Bacteroidetes* (10.5 %), *Planctomycetes* (8.9 %), *Chloroflexi* (7.1 %), *Actinobacteria* (4.7 %), *Verrucomicrobia* (4.7 %), *Acidobacteria* (4.5 %), *Gemmatimonadetes* (4.3 %), *Parcubacteria* (3.3 %) and *Firmicutes* (3.2 %).

To evaluate the overall differences between the microbiomes of the different soil legacies, plant species and plant legacies, we conducted a redundancy analysis (RDA, Oksanen et al., 2017) using the normalized OTU abundances as response variables and the treatment factors with all interactions as explanatory terms (Fig. 2). The two first RDA axes explained 17.4 % of the overall variance and separated the control soil from the monoculture and mixture soils. An exception was “Sample492”, which grouped among the samples from the control soil, even though it came from a microbial soil (see Fig. 2). This sample was therefore removed as outlier from all subsequent analyses. Nonetheless, the result clearly indicated that soils with live inocula had rhizosphere communities that were clearly distinct from the rhizosphere communities that developed in control soil.

**Figure 2.**
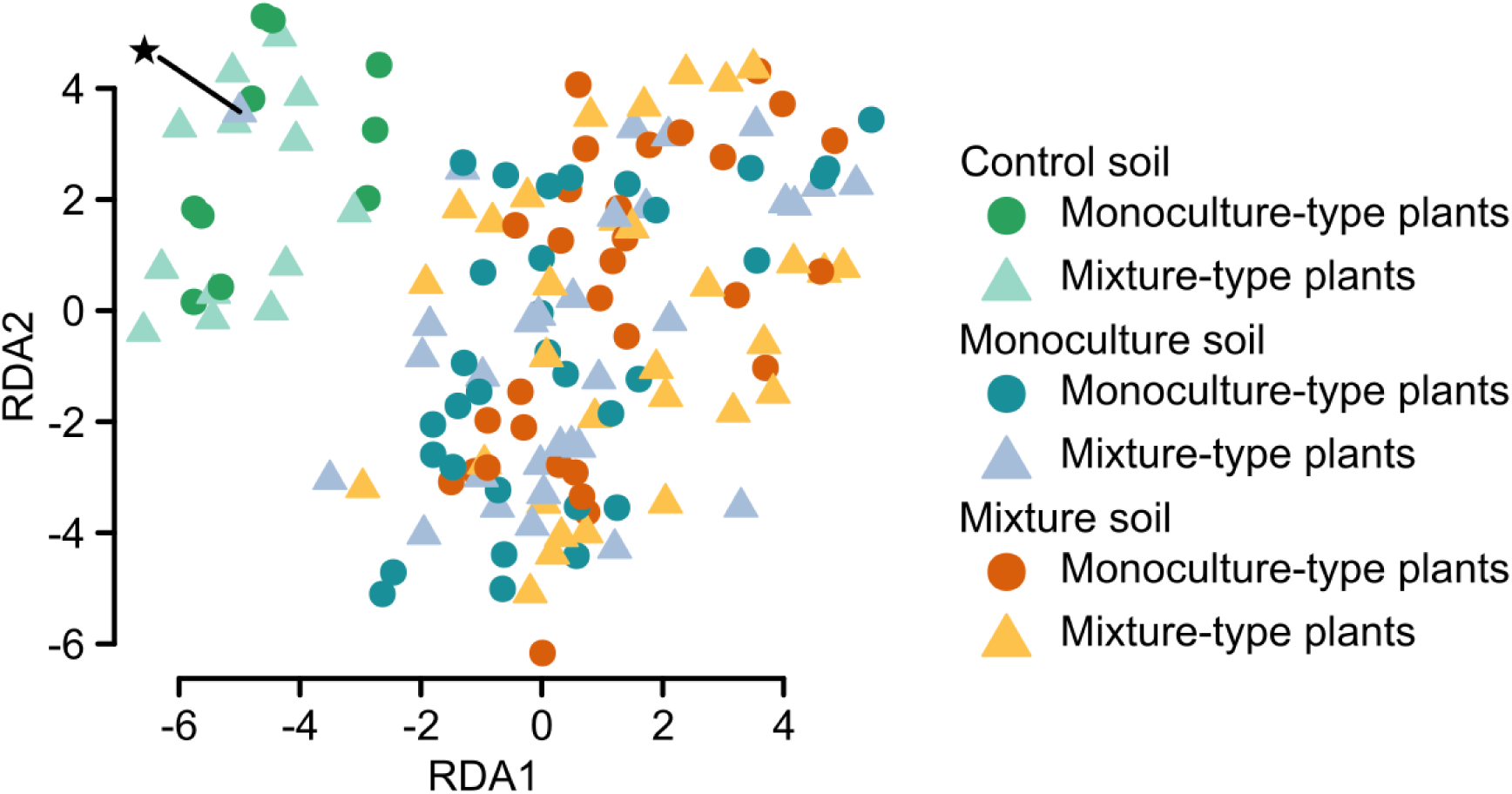
Redundancy analysis using the normalized operational taxonomic unit (OTU) abundances of all samples sequenced. The two first RDA axes explained 17.4 % of the overall variance and separated the control soil from the microbial soils. The constrained components accounted for 60 % of the total variance. The asterisk marks “Sample492” (*L. pratensis*, mixture history, monoculture soil), which clustered among the samples from the control soil. This sample was excluded as outlier from the analysis of differential OTU abundance.

To characterize the overall impact of the treatment factors on microbial diversity, we analyzed the variation in OTU richness among the different treatments (Fig. 3, Table 1). Microbial richness was always higher in the microbial soils than in the control soil, but differences in richness between monoculture and mixture soils were not significant. However, the interactions of these two contrasts with plant species identity were both significant. This indicates that the overall microbial community diversity was primarily determined by the combination of soil-legacy treatments and plant species identity.

**Figure 3.**
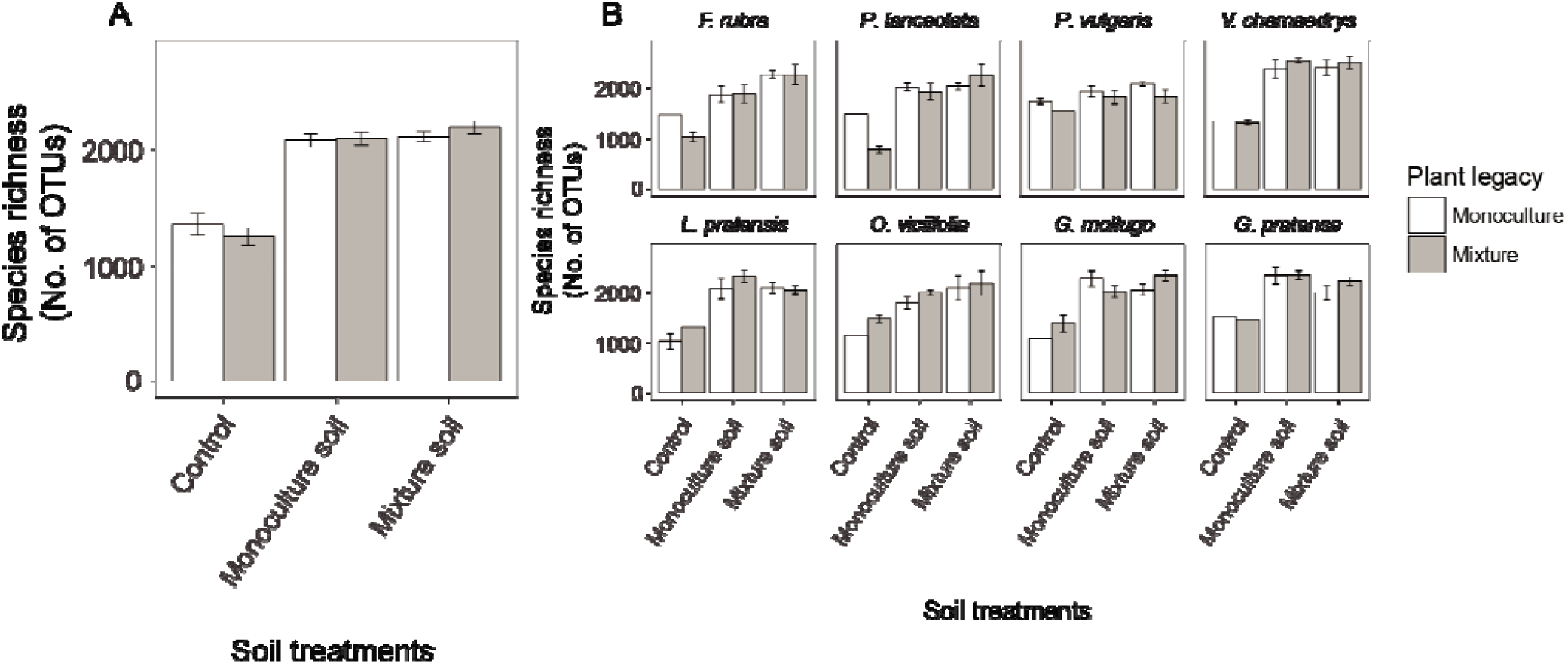
Microbial richness (number of OTUs) in the rhizosphere of monoculture- (white bars) and mixture-type plants (grey bars) across all plant species (A) and for each plant species separately (B). Bars represent means ± standard error.

To identify the OTUs that contributed to the differences in the microbial communities, we tested each OTU for differential abundance between the different soil legacies, plant species identities and plant legacies. We therefore combined the three experimental treatments into a single factor and compared specific conditions with linear contrasts (see Methods for details). Out of the 4’339 OTUs tested, 2’091 showed one or several significant comparisons (Table 2). Comparisons between the different soil-legacy treatments were often significant (e.g., 972 OTUs were different between the control soil and the microbial soils). Likewise, contrasts comparing each plant species to all other plant species were frequently significant (e.g., 498 OTUs were different between *F. rubra* and all other plant species). In agreement with the significant interaction between soil-legacy treatments and plant species identities in the analysis of microbial richness, the number and identity of the OTUs identified as significantly differentially abundant between soil-legacy treatments or among plant species identities varied if they were tested within a given plant species or soil-legacy treatment, respectively (Fig. 4, Fig. S1 and S2). The contrasts comparing the two plant legacies across the entire data set, within microbial soil legacies (monoculture or mixture soil) and within plant species were almost never significant (less than 13 OTUs in every case). Taken together, these results confirmed that the composition of the rhizosphere microbiomes was determined by the microbial community developed over time in the field biodiversity experiment and its interaction with the particular plant species that provided the “root interface” in the pot experiment in the glasshouse. Overall, when looking at the microbial community composition of the normalized abundances of the 2’091 bacterial OTUs which were significant in any of the comparisons, the rhizosphere microbiomes cluster according to plant species and often also according to soil legacy, whereas plant legacies did not form any apparent clusters (Fig. 5).

**Figure 4.**
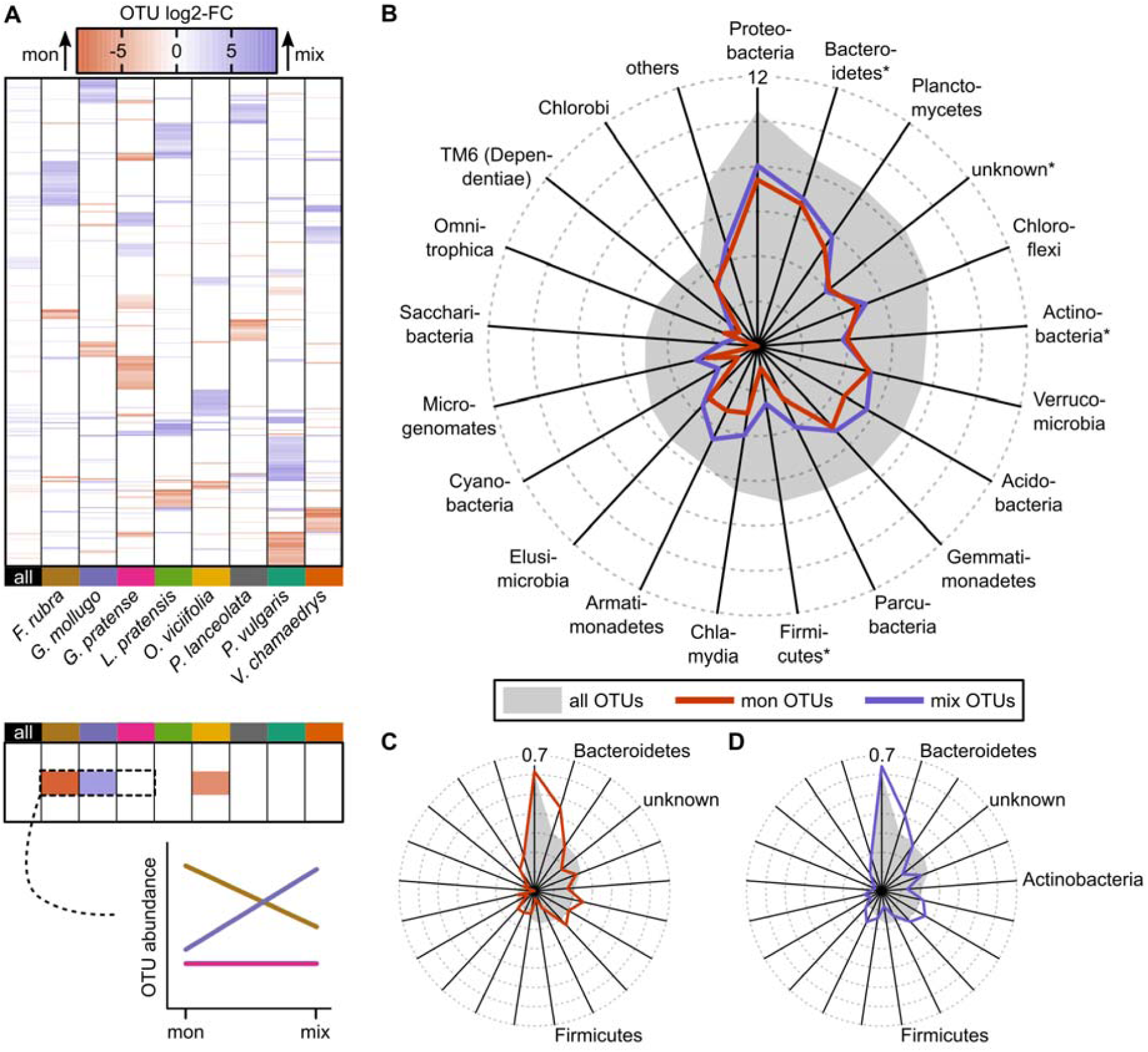
OTUs with significant differences in abundance (log2-FC = log2 fold-changes) between the two soil-legacy treatments monoculture (mon) and mixture soil (mix) across all eight plant species or for each plant species separately (contrast 3. in Table 2). A) Top: Heat map with differences in abundance of the significant OTUs. Each row corresponds to one OTU, each column to the contrast tested across all plant species (“all”) or separately for each plant species. Red or blue corresponds to an increased abundance of an OTU in microbiomes from monoculture or mixture soils, respectively. White indicates an insignificant difference (FDR > 0.01). Bottom: The drawing illustrates how interactions between the plant species and the soil-legacy contrast can be inferred from the heat map. B) OTU frequency (log2-transformed) of the phyla with at least 20 OTUs in the entire data set. The remaining phyla were summarized as “others”. The gray polygon represents the background-distribution (all 4’339 OTUs passing the filters described in Methods). Red and blue lines correspond to the frequencies of OTUs identified as significantly more abundant in microbiomes from monoculture (mon OTUs) and mixture soils (mix OTUs), respectively, within any of the plant species. Phyla with significant enrichment/depletion in either of these sets are marked with asterisks (two-sided Fisher’s exact test, adjusted for multiple testing, FDR < 0.05). C/D) Similar to B), but with OTU frequencies normalized to the total number of OTUs and arc-sin transformed. Only phyla with significant enrichment/depletion are labelled.

**Figure 5.**
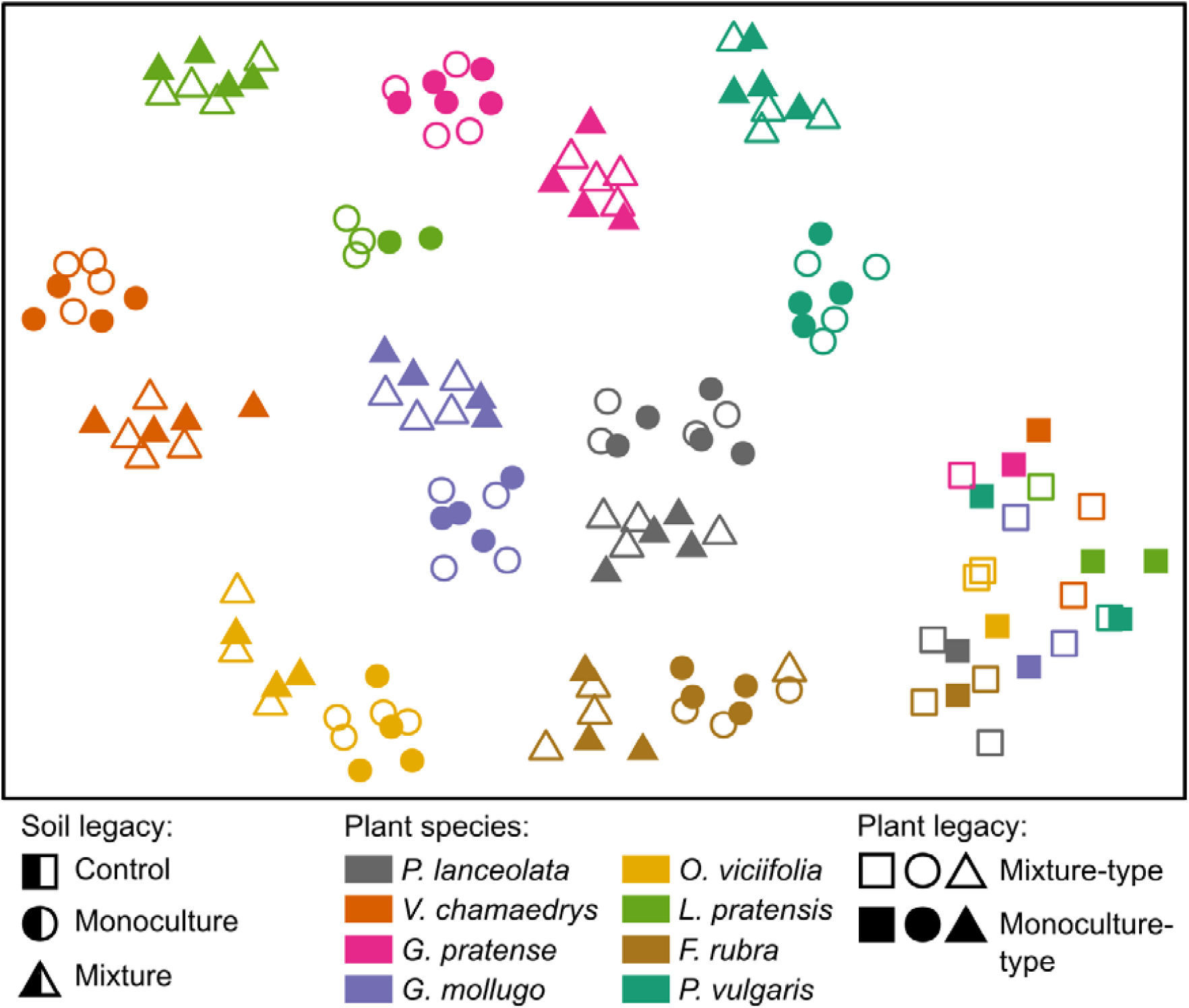
t-SNE map of all samples sequenced and analyzed (excluding the outlier “Sample492”). Control soils (squares) cluster separately. Within plant species (different colors), microbiomes from monoculture soils (circles) and mixture soils (triangles) also cluster separately, however, monoculture- and mixture-type plants within species and soil treatments are associated with similar microbiomes. The map was generated using normalized abundances of OTUs identified as significantly differentially abundant within any of the contrasts tested in this study (2’091 OTUs, Table 2). Note that t-SNE projection axes are arbitrary and dimensions are therefore not shown.

Differences between the microbiomes from monoculture vs. mixture soils were highly specific to each particular plant species (see Fig. 4). In total, 844 OTUs were significantly differentially abundant between the two microbial soils if tested separately for each plant species. The majority of them (566 OTUs) were unique to a given plant species. Only 137 significant OTUs were identified if tested across all plant species (out of which 23 were not among the 844 with significant differences in the plant species-specific comparisons). In contrast, 73.9 % of all OTUs identified as differentially abundant between the control soil and the two microbial soil-legacy treatments were also significant if tested across all plant species (972 out of 1’316 OTUs). On average per plant species, 91 OTUs were more abundant in microbiomes from mixture than from monoculture soil and 64 OTUs were more abundant in microbiomes from monoculture than from mixture soil. Except for *G. pratense*, microbiomes from mixture soil always had a higher number of OTUs with increased abundance than microbiomes from monoculture soil. This was also true if tested across all plant species, where 105 and 32 OTUs exhibited increased abundance in microbiomes from mixture and monoculture soil, respectively (see Table 2). Taken together, the microbiomes from soil with a legacy of plant species mixtures were generally more diverse and contained individual OTUs with higher abundance than the microbiomes from soil with a legacy of monocultures.

We assessed the taxonomy of the OTUs that were significantly differentially abundant between microbiomes from monoculture and mixture soils (Fig. 4B, Table S7). OTUs with increased abundance in microbiomes from monoculture soils were enriched for *Bacteroidetes* (75 observed, 46 expected) and depleted for *Firmicutes* (1 observed, 13 expected) and unknown phyla (14 observed, 36 expected). OTUs with increased abundance in microbiomes from mixture soils were enriched for *Bacteroidetes* (87 observed, 60 expected) and depleted for *Firmicutes* (5 observed, 17 expected) and unknown phyla (11 observed, 47 expected); in addition, they were also depleted for *Actinobacteria* (11 observed, 26 expected).

## 4 DISCUSSION

Here we tested the hypothesis (*i*) that microbiomes obtained from soil from plant mixtures are more diverse and differ in composition from microbiomes obtained from soil from plant monocultures. Secondly, we hypothesized (*ii*) that plant species differ in their microbiomes, and finally, (*iii*) that monoculture-type plants and mixture-type plants differ in their rhizosphere microbiomes when inoculated with microbes from the same soil. Overall, we found distinct differences between control and microbial soil treatments in both microbial composition and richness thus confirming the anticipated establishment of our microbial treatments. Moreover, we were able to show that microbiomes originating from soils from plant monocultures and mixtures did differ, thereby supporting our first hypothesis (*i*), however, this was only apparent in some host plant species. This demonstrates (*ii*) that soil microbiome composition is strongly driven by the host plant species identity directly or in interaction with the soil legacy. However, we found almost no significant differences between microbiomes of monoculture- vs. mixture-type plants, even when testing for interactions with host plant species identity or with soil-legacy treatment and plant species. This provides little support for our hypothesis (*iii*) that plants and their soil microbiome have been co-selected. However, it is also possible that co-selection of microbes occurred at a taxonomic level for which the resolution of 16S rRNA sequencing is too low (e.g., if plants selected different strains of the same microbial species).

Taken together our findings demonstrate the composition of the soil microbiome is shaped by the environment from which they originated in which plant diversity legacy effects play a significant role, and by the identity of the host plant species with which they associate during plant growth.

### Effects of microbiome legacy

It is important to note that the mixture and monoculture soil inocula included microbial filtrates from eight different plant monoculture plots and seven different eight-species plant mixture plots (one for each species except for *G. mollugo* and *O. viciifolia* whose rhizosphere soil samples came from the same mixture plot) in the Jena Experiment. To a certain extent, the average monoculture soil thus represents a mixture influenced by eight plant species. Likewise, the average mixture soil represents a mixture influenced by seven different plant communities (36 plant species in total). The overall comparison between monoculture and mixture soil across all plant species thus resembles a comparison of soil influenced by eight plant species compared to 36 plant species (without considering the potential effects of plant diversity on each individual soil sample from the Jena Experiment). The effect of soil legacy on the microbial diversity may therefore be better understood when comparisons are made within each plant species.

### Factors affecting microbiome diversity

Indeed, differences in bacterial OTU richness between microbiomes from monoculture and mixture soil varied among the eight plant species (see Fig. 3B and Table 1). Considering that plant species identity was confounded with species composition of plant mixtures in the Jena Experiment, it is possible that in part these differences were due to key plant species being present or absent in these plant mixtures in the Jena Experiment. However, a more diverse plant community always has a higher chance to contain some particular species. In this sense, the presence of a key species may also be considered as a biodiversity effect, sometimes referred to as “selection probability effect” (Niklaus, Baruffol, He, Ma, & Schmid, 2017). Only three of the eight studied plant species showed clearly increased bacterial richness in microbiomes from mixture than from monoculture soil when the total numbers of detected OTUs were analysed.

However, bacterial richness is unlikely a robust measure to compare different conditions because it does not take into account the sequencing depth and more importantly, the differences between bacterial abundances. Compared to species abundances of larger organisms (e.g. flowering plants), abundances of bacterial species are estimated less reliably because sequencing of 16S rRNA provides only an indirect abundance estimate, which can be influenced by numerous technical factors. Also, bacterial abundance can vary much more than abundances of larger organisms. For example, using the normalized read counts as a proxy for abundance, the “abundance” of all OTUs in the data presented ranged from 0 to 17’420. Likewise, the differences in abundance between conditions reached up to a log2-fold change of 9.72 (a factor of 843 on linear scale). Finally, read counts may originate from leftover DNA in the soil instead of living bacteria. Thus, choosing a particular threshold to define presence/absence of a bacterial species may result in different interpretations. Furthermore, different thresholds may be appropriate to indicate presence/abundance in different OTUs (Edgar 2017). To overcome these limitations of bacterial species richness based on presence/absence data, we focused on the results of the analysis of differential abundances of given bacterial OTUS.

### Factors affecting microbiome abundance

When we compared OTU abundances between the microbiomes from mixture and from monoculture soils, the number of OTUs with increased abundance in mixture soil was higher in all plant species with the exception of *G. pratense* (see Fig. 4 and Table 2). These results are in line with previous studies reporting a positive correlation between plant diversity and soil bacterial abundances (Eisenhauer et al., 2017; Stephan, Meyer, & Schmid, 2000), or generally soil microbial abundances (Eisenhauer et al., 2013; Thakur et al., 2015). Studies examining the correlation of plant species diversity and soil bacterial richness showed positive correlations (Garbeva et al., 2006; Stephan et al., 2000), negative correlations (Schlatter et al., 2015) or no correlation (Dassen et al., 2017). Because the habitats and resources in the rhizosphere tend to vary between different plant species (Berg & Smalla, 2009; Eisenhauer et al., 2017; Hooper et al., 2000), increasing plant species diversity could provide larger variety of resources and habitats for microbes and thereby explain the higher bacterial abundances of several taxa and higher total richness of microbiomes from mixture than from monoculture soil observed here. Additionally, the clear differences in bacterial composition between the microbiomes from the two soil legacies and the weak influence of plant legacies in the eight plant species (see Fig. 5) indicate that the differences in bacterial abundance and richness did not develop much during the five months of the pot experiment in the glasshouse, but were primarily determined by the 11 (=8+3) years of co-development between plants and soil microbial communities in the field plots of the Jena Experiment.

Only in the tall herb *G. pratense* bacterial abundance and richness was larger in microbiomes from monoculture than from mixture soil. One explanation could be the selection history of *G. pratense*. Whereas all the other species grew in mixed-species field plots with herbs and grasses, legumes or both, *G. pratense* was growing in a mono-functional group mixture (eight species of tall herbs) in the Jena Experiment. This may have resulted in a more monoculture-like selective environment for the associated microbes. For the other plant species growing in species mixtures with more than one functional group, bacterial abundance and richness were consequently larger for microbiomes from mixture than from monoculture soil. Plant functional groups have been shown to influence bacterial abundance (Stephan *et al*., 2000; Bartelt-Ryser *et al*., 2005; Latz *et al*., 2012, 2016; Lange *et al*., 2014) and richness (Stephan *et al*., 2000; Dassen *et al*., 2017) in soil. In the last study, Dassen et al. (2017), suggested that plant functional groups are more important determinants of bacterial richness than plant species. Taken together, our results suggest that bacterial abundance and richness in the rhizosphere generally increase with increasing plant species diversity to the extent that this feeds back to newly establishing plants, but that they are also positively influenced by plant functional diversity.

## 5 CONCLUSIONS

Our results suggest that plant diversity generally increased the diversity of soil bacteria in the rhizospheres of eight plant species. The exception of *G. pratense* suggests that this effect can also depend on the identity of the host plant species. These findings support our hypothesis that when plants and soil microbial communities are allowed to develop together for prolonged time periods in plant monocultures and mixtures in the field, the diversity and composition of bacterial communities subsequently associated with plant roots can diverge. High biodiversity both above- and belowground may provide an important insurance for plant biomass production in the long term. Furthermore, these results emphasize that concerns about plant biodiversity loss that may also have cascading effects on to soil biodiversity loss and functioning of terrestrial ecosystem processes.

## ACKNOWLEDGEMENTS

Thanks to D. Flynn for help with the experimental design. We furthermore thank M. Furler, H. Martens, D. Topalovic, D. Trujillo Villegas and T. Zwimpfer for technical assistance, M. Brezzi, Y. Xu, E. De Luca and A. Grossniklaus for help with the experiment and Agroscope for providing facilities. This study was supported by the Swiss National Science Foundation (grants number 147092 and 166457 to B. Schmid) and the University Research Priority Program Global Change and Biodiversity of the University of Zurich. The Jena Experiment is supported by the German Science Foundation (FOR 1451, SCHM 1628/5-2). G. B. De Deyn was financially supported by NWO-ALW VIDI (grant number 864.11.003).

## DATA ACCESSIBILITY

Short-reads were deposited at SRA (accession number SRP105254). Supplementary tables and files are accessible on https://dx.doi.org/10.5281/zenodo.1044377.

## AUTHOR CONTRIBUTION

T.H., C.W. and B.M. designed the study. T.H. and S.J.V.M. carried out the experiment. T.H. performed the DNA extraction and sequencing preparation. M.W.S. processed the sequencing data and performed data analysis. The paper was written by S.J.V.M, M.W.S. and T.H. with all authors contributing to the final version.

