## Supplementary Materials for "Rhizosphere bacterial community composition depends on plant diversity legacy in soil and plant species identity"

**Supporting Information**

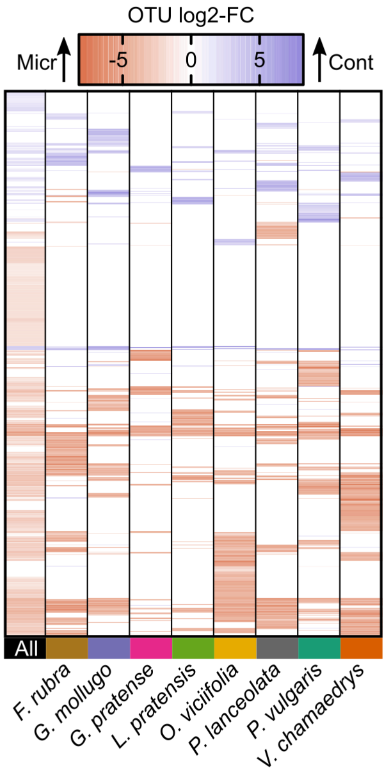


**Figure S1.** OTUs with significant differences in abundance (log2-FC = log2 fold-changes) between the the control (cont) and microbial soil (micr) (contrast 2. in Table 2). Heatmap with differences in abundance of the significant OTUs. Each row corresponds to one OTU, each column to the contrast tested across all plant species ("all") or separately for each plant species. Blue or red corresponds to an increased abundance of an OTU in microbiomes from control or microbial soils, respectively. White indicates an insignificant difference (FDR > 0.01).

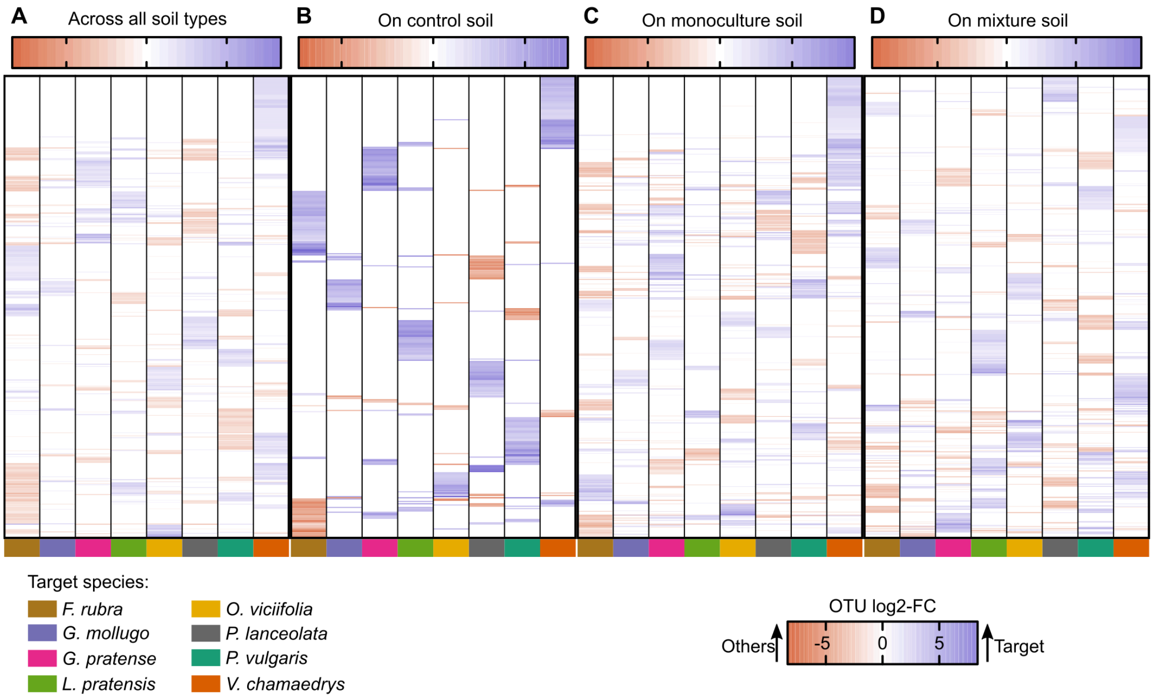


**Figure S2.** OTUs with significant differences in abundance (log2-FC = log2 fold-changes) between a given plant species (target) and all other plant species (others) across all soil-legacy treatments (A, contrast 4. in Table 2) or for each soil-leagy treatment separately (B–D, contrasts 4.a)–4.c) in Table 2). Heatmap with differences in abundance of the significant OTUs. Each row corresponds to one OTU, each column to the comparison of one selected plant species with the average of all others. Blue or red corresponds to an increased abundance of an OTU in microbiomes from the target or all other plant species, respectively. White indicates an insignificant difference (FDR > 0.01).

**Supporting Files (accessible on Zenodo)**

**Table S1.** The table contains the annotation for all the samples sequenced and analyzed. Accessible on https://dx.doi.org/10.5281/zenodo.1044377.

**Table S2.** The table contains all primer sequences used in this study. Accessible on https://dx.doi.org/10.5281/zenodo.1044377.

**Table S3. (OTU_annotations.csv.zip)** The zip-file contains a table with the taxonomic annotation of the operational taxonomic units (OTUs) identified in this study. Accessible on https://dx.doi.org/10.5281/zenodo.1044377.

**Table S4. (OTU_counts.csv.zip)** The zip-file contains a table with sequence counts of the operational taxonomic units (OTUs) identified in this study. Accessible on https://dx.doi.org/10.5281/zenodo.1044377.

**Table S5. (OTU_diffAbundance.xlsx)** The workbook contains a sheet with the number of operational taxonomic units (OTUs) exhibiting differential abundance in any of the contrasts tested in this study. Note that ”down/up“ indicates whether the OTU was less (down) or more (up) abundant in the first group of the contrast. For example, given the contrast “PH_mix_vs_mon_”, down corresponds to higher abundance in the pots with monoculture- than the pots with mixture-type plants. Conversely, up refers to higher abundance in the pots with mixture-type plants. In addition, the workbook contains one sheet per contrast with the logBaseMean (log2 of the average normalized abundance across all samples), the logFC (log2 of the fold-change), the *P*-value, and the adjusted *P*-value (FDR). Only OTUs with a *P*-value <= 0.05 or an adjusted *P*-value <= 0.1 are given. Accessible on https://dx.doi.org/10.5281/zenodo.1044377.

**Table S6. (OTU_overallTaxonomy.csv)** The table contains the number of bacterial OTUs annotated with a given bacterial phylum. Accessible on https://dx.doi.org/10.5281/zenodo.1044377.

**Table S7 (phylaEnrichment.csv)** The table contains all phyla tested for enrichment/depletion in the set of OTUs with an increased abundance in monoculture and mixture soil, respectively. “Total counts (all OTUs)” corresponds to the total number of all OTUs annotated with a given phyla (reference set). “Observed” corresponds to the number of OTUs annotated with a given phylum in the set of OTUs with increased abundance in monoculture or mixture soil (test set). “Expected” gives the number of OTUs which would be expected to be annotated with a given phylum if the test sets were randomly sampled from the reference set. Accessible on https://dx.doi.org/10.5281/zenodo.1044377.

**File S1. (OTU_references.fasta.zip)** The zip-file contains a fasta file with the 10'205 OTU sequences identified in this study. Accessible on https://dx.doi.org/10.5281/zenodo.1044377.
